# Estimation of cancer cell fractions and clone trees from multi-region sequencing of tumors

**DOI:** 10.1101/2021.06.12.448194

**Authors:** Lily Zheng, Laura Wood, Rachel Karchin, Robert Scharpf

**Affiliations:** Department of Genetic Medicine Institute for Computational Medicine Johns Hopkins University School of Medicine Baltimore, MD 21205; Department of Oncology Department of Pathology Johns Hopkins University School of Medicine Baltimore, MD 21287; Department of Biomedical Engineering Institute for Computational Medicine Department of Oncology Johns Hopkins University and Medicine Baltimore, MD 21218; Department of Oncology Johns Hopkins University School of Medicine Baltimorex, MD 21287

**Keywords:** clone trees, Bayesian inference, tumor evolution, next generation sequencing, pancreatic cancer, graphical models, copy number, somatic mutations

## Abstract

Multi-region sequencing of one or multiple biopsies of solid tumors from a patient can be used to improve our understanding of the diversity of subclones in the patient’s tumor and shed light on the evolutionary history of the disease. Due to the large number of possible evolutionary relationships between clones and the fundamental uncertainty of the mutational composition of subclones, elucidating the most probable evolutionary relationships poses statistical and computational challenges. We developed a Bayesian hierarchical model called PICTograph to model uncertainty in the assignment of mutations to subclones and an approach to reduce the space of possible graphical models that postulate their evolutionary origin. Compared to available methods, our approach provided more consistent and accurate estimates of cancer cell fractions and better tree topology reconstruction over a range of simulated clonal diversity. Application of PICTograph to whole exome sequencing data of individuals with pancreatic cancer precursor lesions confirmed known early occurring mutations and indicated substantial molecular diversity, including multiple distinct subclones (range 6 - 12) and intra-sample mixing of subclones. As the complete evolutionary history for some patients was not identifiable, we used ensemble-based visualizations to distinguish between highly probable evolutionary relationships recovered in multiple models from uncertain relationships occurring in a small subset of models. These analyses indicate that PICTograph provides a useful approximation to evolutionary inference, particularly when the evolutionary course of a patient’s cancer is complex.

## Introduction

Multi-region and multi-lesion tumor sequencing studies have motivated development of statistical methods that infer the evolutionary history of tumors by modeling latent processes of clonal expansion and selection. Methods for modeling cancer evolutionary trees from multi-region sequencing follow two main strategies: sample trees and clone trees. In a sample tree, each sequenced tumor is a leaf node and internal nodes represent unobserved ancestral states [Gerlinger et al., 2014, Ling et al., 2015, Zhao et al., 2016, Zhai et al., 2017]. An implicit assumption of sample tree analyses is that the samples are monoclonal or can be meaningfully summarized as the collection of observed mutations, and thus reflect the overall similarity of samples [Alves et al., 2017]. This assumption can lead to sample trees that give the appearance of a somatic variant occurring independently on different branches when the more probable phenomenon is the presence of multiple subclones, some of which are shared between the sequenced regions. The inability to represent intratumor heterogeneity and the evolutionary relationships among different tumor subclones has lead to a rapidly developing focus in the community on clone tree models where each node represents a cluster of mutations, and each extant population of cells comprising a subclone is represented by the aggregation of mutations from root to leaf [Niknafs et al., 2015, Deshwar et al., 2015, Popic et al., 2015, Jiang et al., 2016, Satas and Raphael, 2017]. Identifying clusters of mutations likely to have arisen at the same time and ordering mutation clusters in a branching tree topology are among the main computational challenges of clonal tree inference from multi-region sequencing.

Clone tree models rely on an estimate of the proportion of cancer cells in a tumor that harbor a somatic mutation (cancer cell fraction, CCF), as well as the number of subclones. Unlike the variant allele fraction (VAF) where direct estimates are available from standard mutation-calling algorithms [Benjamin et al., 2019], the CCF is not observed directly and must be inferred from the VAF, multiplicity, DNA copy number of the tumor genome containing the mutation, and the tumor purity of the bulk sample that was sequenced. As heterogeneity in the mutational composition of subclones give rise to differences in the observed VAF between mutations, statistical approaches for clustering mutations provide an avenue to inferring the number of subclones and improving CCF estimation by pooling information from all available mutations within a subclone [Roth et al., 2014, Miller et al., 2014, Niknafs et al., 2015]. Bayesian mixture models implemented by Markov Chain Monte Carlo (MCMC) [Roth et al., 2014] or variational Bayes [Miller et al., 2014, Gillis and Roth, 2020] are attractive for their ability to jointly summarize the uncertainty of CCF estimates and the assignment of mutations to clusters through posterior distributions. These approaches for clustering mutations differ by the types of mutations analyzed (sequence-only or copy number and sequence), the determination of the number of clusters, and how unknown parameters including the VAF, copy number, multiplicity, and purity are modeled to infer CCFs.

Mutation clusters characterized by their CCFs can be ordered in a branching tree topology [Deshwar et al., 2015, Niknafs et al., 2015, Jiang et al., 2016, Satas and Raphael, 2017]. These topologies are generally restricted by the Sum Condition [Niknafs et al., 2015, Popic et al., 2015, El-Kebir et al., 2015, Satas and Raphael, 2017], which states that the CCF of an ancestral clone must be greater than or equal to the sum of CCFs of its descendants. This principle can be applied to pairwise comparisons as the lineage precedence rule [Niknafs et al., 2015], which states that the CCF of any mutation cannot exceed the CCF of its ancestor. Existing tree inference methods apply these principles using probabilistic [Niknafs et al., 2015, Deshwar et al., 2015, Jiang et al., 2016] and combinatorial [Popic et al., 2015, El-Kebir et al., 2015, Satas and Raphael, 2017] frameworks. Due to the number of somatic mutations and likely subclones in many solid tumors, efficient approaches to explore the most probable trees are needed.

We developed a Bayesian hierarchical model called PICTograph to cluster mutations by posterior summaries of CCFs and develop simple assumptions based on the probable absence of mutations in a sample that greatly simplify the space of possible trees. Using simulated data, we evaluate PICTograph when the true clonal diversity and evolutionary history of mutations is known and compare PICTograph to current state-of-the-art clone tree methods. We then applied PICTograph to multi-region whole-exome sequencing (WES) of pancreatic cancer precursor lesions from Fujikura *et al*. [Fujikura et al., 2020] that were previously analyzed using a sample tree approach [Reiter et al., 2017], highlighting beneficial insights aided by the clone tree model. Through this effort, we provide a statistical framework and software for clone-tree inference of multi-region sequencing data obtained from patients with cancer (https://github.com/karchinlab/pictograph).

## Results

### Overview of Approach

We developed a generative model for observing somatic mutations in read-level data from whole exome sequencing with unknown parameters that include the VAF, the CCF of the clone, and the number of clones (Figure 1). As patients with as few as 10 estimated mutation clusters, *K*, would have more than 2 billion possible evolutionary relationships ((*K* + 1)^*K*−1^ or 11^9^ trees [Cayley, 1889]), exhaustive computational methods to evaluate the number of possible trees are untenable. To overcome this limitation and restrict inference to more probable trees, our approach groups mutations by the set of samples in which the mutations are detected (sample presence, Figure 1) [Popic et al., 2015, Myers et al., 2019]. CCF estimation and cluster assignments are performed independently within sample presence strata, establishing a clone tree hierarchy in the mutation groups that can reduce the number of possible trees by orders of magnitude. We adopted a hierarchical Bayesian model for the CCFs such that the posteriors integrate information from all available mutations assigned to the same subclone. Using the posterior modes of the CCFs, we determined the collection of possible trees consistent with the sample presence hierarchy and ranked the trees using a score that quantifies the compatibility between the tree and CCF estimates. To evaluate and benchmark this approach, we simulated clone trees capturing a broad range of complexity and implemented multiple currently available clone-tree methods. Finally, we applied PICTograph to publicly available experimental data where multi-region whole exome sequencing was performed to characterize the clonal composition of somatic mutations in pancreatic cancer precursor lesions [Fujikura et al., 2020].

**Figure 1:**
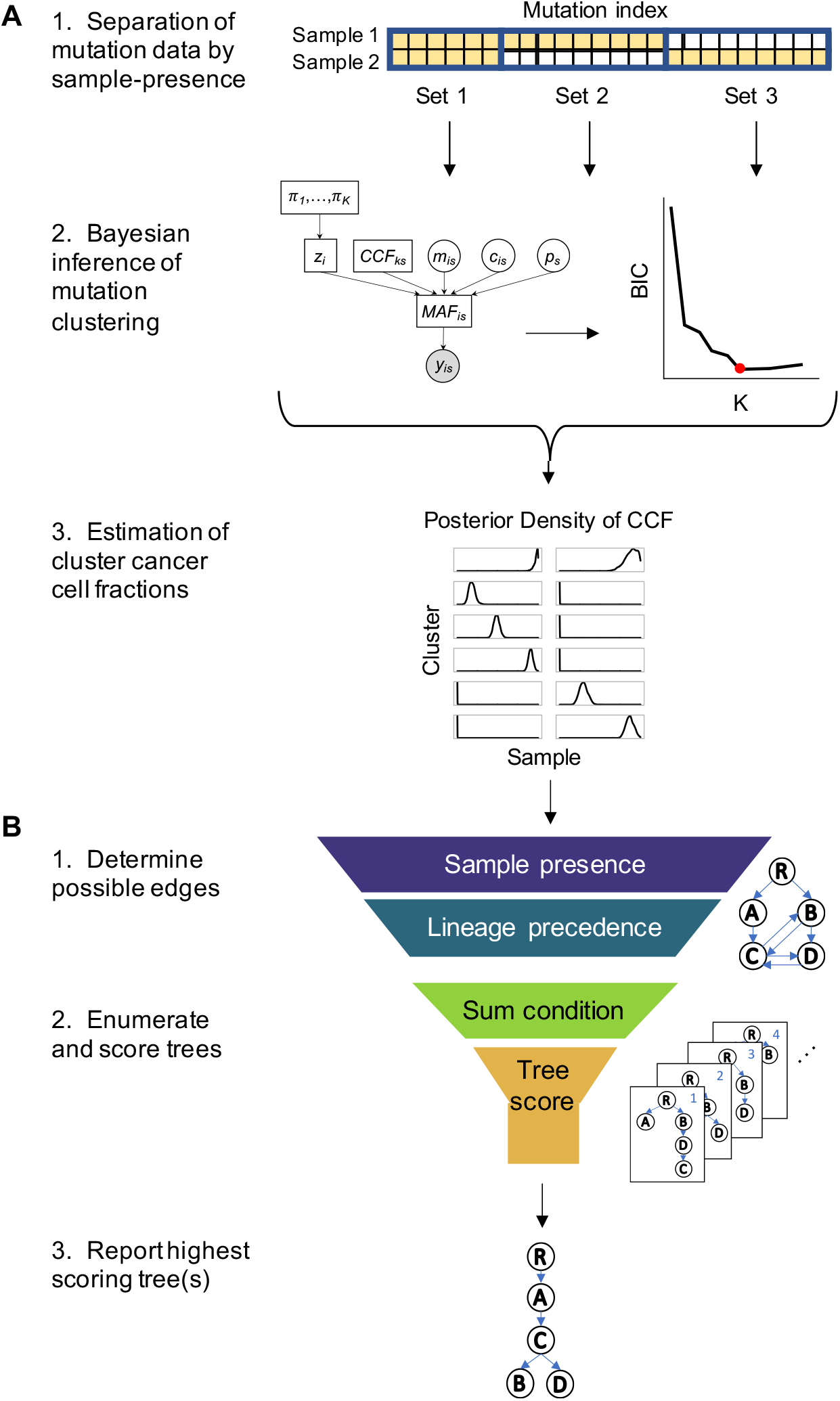
Overview of approach for processing multi-sample next generation sequencing data. The PICTograph algorithm contains two main components: CCF estimation and tree inference. (A) To estimate CCFs, PICTograph first separates the mutation data into sets based on sample presence patterns. Then for each mutation set, PICTograph uses MCMC sampling to jointly estimate each mutation’s cluster assignment and the CCFs of each cluster for a plausible range of mutation clusters *K*. The Bayesian Information Criterion (BIC) is applied to select the optimal value of *K*. The total number of clusters of a case is the sum of the number of clusters identified in each set. PICTograph calculates maximum a posteriori estimates of the cluster CCFs to use for subsequent tree inference. (B) For tree inference, PICTograph uses estimates of cluster CCFs to determine a set of possible edges by applying filters based on sample presence and the lineage precedence rule. From this set of edges, PICTograph uses the Gabow-Myers algorithm to enumerate all spanning trees, and then further filters trees based on the Sum Condition. Finally, this set of filtered trees is scored with the SCHISM fitness function, and the highest scoring trees are reported and summarized as an ensemble tree.

### Application of PICTograph to multi-region whole exome sequencing of tumors

Following the alignment and identification of somatic mutations in each sequenced lesion of a patient, our approach stratifies the identified mutations into tiered sets by sample-presence. The first tier consists of mutations present in all of the samples and consists of a single set. The second tier is formed by mutations present in all but one of the samples and consists of at most 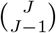 sets. Mutations in lower tiers are assigned to sample sets in a similar manner until all mutations have a sample set designation (Figure 1A). Independently for each sample-presence tier, we modeled the observed allele frequency for a mutation in a sample using a binomial sampling distribution parameterized by the VAF (Methods). An identity link function decomposes the VAF into the CCF of the clone to which the mutation belongs, accounting for the mutation-specific copy number, multiplicity, and sample-specific estimates of tumor purity. As the mutational composition of the subclones in a sample are themselves unknown, we introduce an auxillary variable that assigns the mutation to a subclone and a prior probability distribution for this latent variable, as is standard in Bayesian mixture models. As the CCF of a tumor subclone would generally contain multiple mutations before it could be reliably distinguished from other subclones, we model the CCFs hierarchically so that information about the CCF of a clone is informed by all available mutations assigned to the clone. PICTograph approximates the joint posterior distribution of the mixture component membership and CCF by Markov Chain Monte Carlo (MCMC).

To determine the likely evolutionary relationships consistent with sample presence and lineage precedence, we first summarized the CCF and mutation cluster assignment by the posterior modes. Representing the set of allowed relationships (edges) between mutation clusters (nodes) in a directed acyclic graph, we used the Gabow-Myers algorithm [Gabow and Myers, 1978] to enumerate all possible trees consistent with lineage precedence and the Sum Condition. As these requirements could potentially exclude the true tree, we employed soft filters on both the lineage precedence and Sum Condition (Methods) and scored trees using SCHISM [Niknafs et al., 2015] to penalize any violations. PICTograph reports the highest scoring tree and, in the event of a tie, summarizes the collection of trees with the maximum score as an ensemble tree (e.g., Figure 1).

### Simulations and benchmarks against state-of-the-art methods

We first evaluated our approach through simulation of multi-region tumor sequencing data simulated from known clone trees and CCF values for each mutation cluster. We generated 100 random trees each for 5, 10, or 15 mutation clusters. For each case, we randomly simulated 3 tumor samples with tumor purity ranging from 50 to 90%, allocated 1 to 10 mutations per cluster (e.g., Figure 2A), and generated compatible CCFs from a uniform Dirichlet distribution [Dentro et al., 2017]. Using the generative model (Figure 1A), we simulated read level data of 100x average coverage.

**Figure 2:**
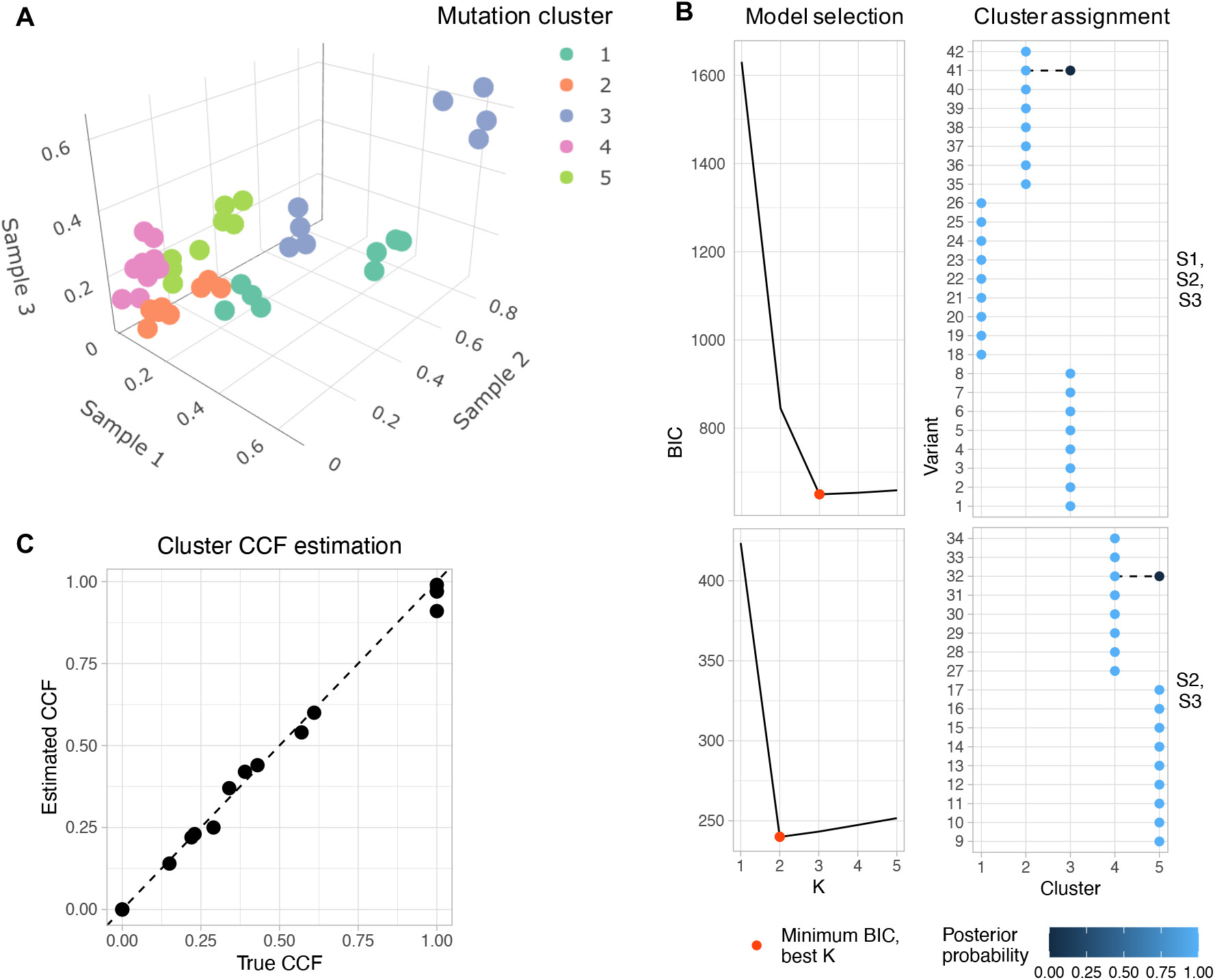
Clustering performance of PICTograph on simulated data. (A) Multidimensional scatter plot of variant allele frequencies (VAFs) for all mutations across 3 samples shows groupings of mutations in 3D space. Colors represent mutation cluster assignments by PICTograph, which are also in agreement with the true cluster assignments. (B) For each sample-presence set, PICTograph performs model selection and inference of mutation clusters. (C) Maximum a posteriori estimates of cluster CCFs are nearly identical to the true CCFs and are used for subsequent tree inference.

**Figure 3:**
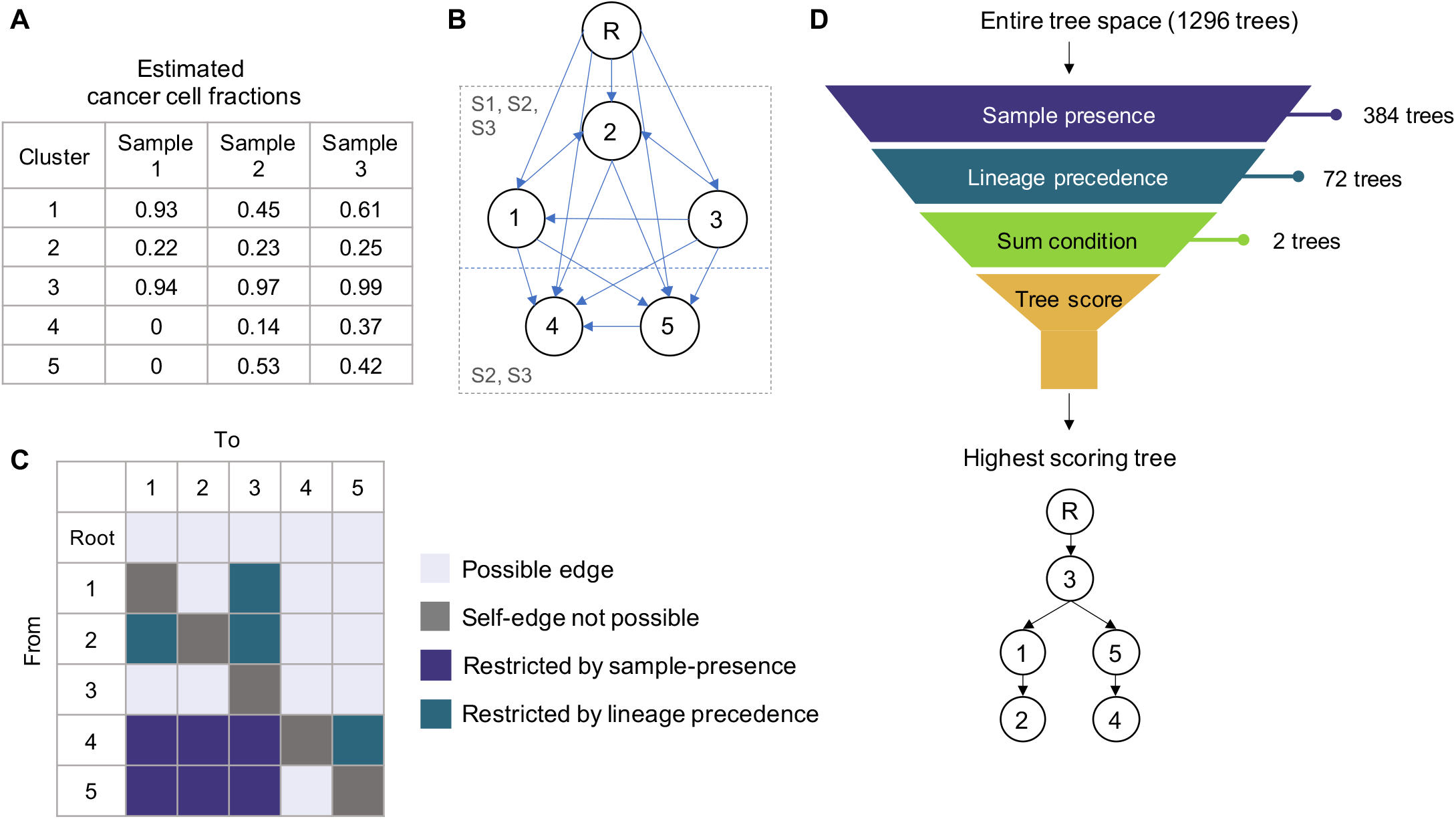
Tree inference of simulated data using PICTograph. (A) PICTograph summarizes the CCF posterior distribution for each cluster by the mode. (B) Mutation clusters (numbered circles) are shown in boxes representing their respective sample-presence sets. The root node is shown as a circle labeled R located above the sample-presence sets. Arrows represent the possible edges determined by applying filtering rules to pair-wise comparisons of CCF estimates of mutation clusters. (C) An adjacency matrix representation of possible edges and those restricted by sample-presence and lineage precedence filters. (D) We use sample presence, lineage precedence, and the sum c ondition to reduce the tree space. A tree scoring function that penalizes violations of lineage divergence and precedence is used to report the highest scoring tree(s).

For each simulation, we estimated the number of mutation clusters, the cancer cell fractions of the mutation clusters in each sample, and scored trees consistent with the estimated cancer cell fractions as previously described. To benchmark our approach against current state-of-the-art methods, we implemented approaches for mutation clustering and CCF estimation (PyClone [Roth et al., 2014] and sciClone [Miller et al., 2014]), tree inference (LICHeE [Popic et al., 2015]), and methods that infer both mutation clusters and clone trees (Canopy, PhyloWGS, SCHISM). We evaluated methods by their ability to correctly identify the true number of mutation clusters, the average distance between CCF estimates and true CCF, and compared mutation trees by the proportion of correctly identified evolutionary relationships. As expected, the performance of all methods decreased as the true number of clusters increased. PICTograph and Canopy had the lowest level of error for inferring the correct number of mutation clusters with a mean absolute errors of 2.69 (IQR = 1-4) and 2.89 (IQR = 1-5), respectively, compared to SCHISM, PhyloWGS, sciClone, and PyClone that had mean absolute errors of 5.58 (IQR = 1-9), 8.17 (IQR = 3-13), 5.16 (IQR = 2-8), and 4.32 (IQR = 2-7), respectively (Figure 2A). On average, PICTograph, SCHISM, PhyloWGS, sciClone, and PyClone underestimated the number of clusters while Canopy overestimated the number of clusters. Next, we measured accuracy of the CCF estimates using two complimentary metrics of divergence (see Methods). Comparing the divergence of these metrics for each method to PICTograph, we calculated the proportion of simulated datasets with lower divergence by PICTograph for Metrics 1 (*p*_1_) and 2 (*p*_2_). PICTograph had lower divergence than PhyloWGS, PyClone and sciClone regardless of the number of true clusters (*p*_1_ = 1, Figure 4). PICTograph and Canopy performed similarly for simulations with only 5 clusters (*p*_1_ = 0.53) but PICTograph had lower measures of divergence for simulations with 10 (*p*_1_ = 0.72) and 15 clusters (*p*_1_ = 0.86). Similarly, SCHISM performed well for simulations with few clusters (*p*_1_ = 0.41) but performance declined with increasing complexity of the simulations (*p*_1_ = 0.79 for 10 clusters and *p*_1_ = 0.91 for 15 clusters). With respect to Metric 2, we found that PICTograph had lower measures of divergence regardless of cluster size for both Canopy (*p*_2_ ≥ 0.92) and PyClone (*p*_2_ range: 0.84-0.97). Both PhyloWGS and SCHISM had lower measures of divergence for Metric 2 (*p*_2_ range: -0.63 - 0.50 for PhyloWGS and -0.62-0.23 for SCHISM), but the lower difference by this metric was achieved by significantly underestimating *K* with much higher divergence by Metric 1 (Figures 4A and B). Overall, these results indicate that PICTograph provided more accurate estimates of CCF and *K* than other methods across a broad range of simulated subclonal complexity.

**Figure 4:**
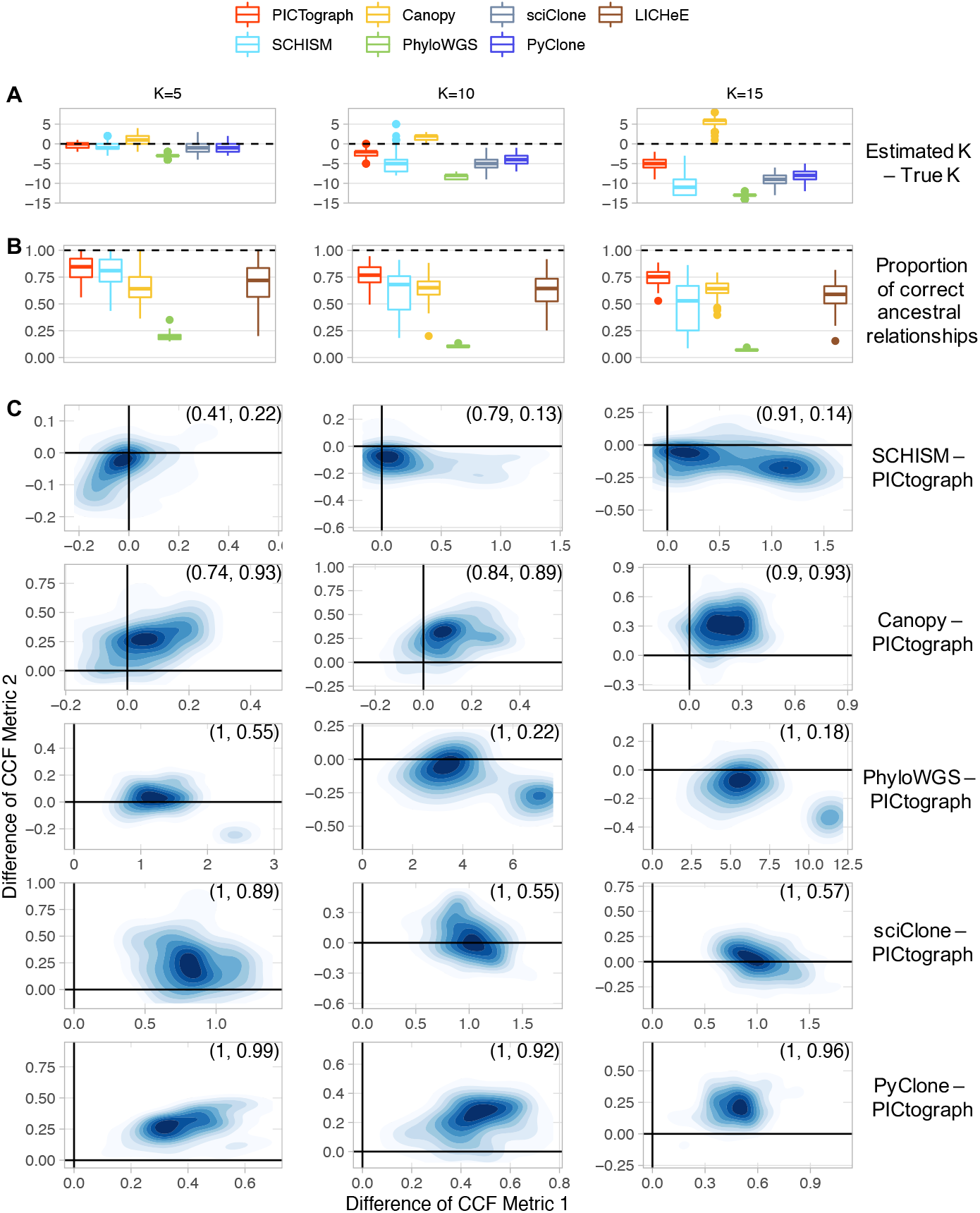
Comparison of evolutionary methods. We simulated 100 cases each for 5, 10, and 15 vertex (*K*) trees. For each method, we calculated the difference between the estimated number of clusters and the true number of clusters (A) and the proportion of ancestral relationships correctly inferred (B). Dotted lines indicate the best possible score. (C) To compare the distribution of estimated CCFs to the true CCF distribution, we used two assymetric measures of divergence that penalize underfitting (Metric 1) or overfitting (Metric 2). Comparing each divergence metric to PICTograph, data points in the top right quadrant indicate lower divergence by PICTograph. The proportion of simulations for which the divergence was smaller in PICTograph by Metrics 1 and 2 are indicated in the top right panel as (*p*_1_, *p*_2_). Collectively, these simulations indicate that PICTograph performs better than alternative methods particularly as the complexity of the simulated data increases with more mutation clusters. Methods that perform well by CCF metric 2 (SCHISM and PhyloWGS) in comparison to PICTograph tend to greatly under-estimate the true number of clusters (corroborated by panel A) and perform poorly by Metric 1.

Next, we evaluated the ability of PICTograph, SCHISM, Canopy, PhyloWGS, and LICHeE to identify subclone trees. Ranking trees by their score (SCHISM, PICTograph, and LICHeE) or posterior probability (Canopy), we calculated accuracy as the proportion of correctly inferred ancestral relationships from all pairwise mutation comparisons [Satas and Raphael, 2017] for the top-ranked tree or the average accuracy in the event of tied trees. Since PhyloWGS provides a posterior distribution of clone trees, we calculated the mean posterior accuracy (Figure 4D). Overall, we found that PICTograph had the highest average accuracy (mean = 78.2%, IQR = 71.2-85.5%). Accuracies were lower for SCHISM (mean = 63.2%, IQR = 50.2-79%), Canopy (mean = 64.5%, IQR = 58.4-70.8%), and LICHeE (mean = 63%, IQR = 52.9-75.3%). PhyloWGS had the worst accuracy (mean = 12.3%, IQR = 7.5-10.3%), likely due to poor clustering and its effect on the measured accuracy of evolutionary relationships (Figure 4D). Collectively, these simulations indicate that recovery of ancestral relationships between subclones by PICTograph is robust to a wide range of mutation clusters and tree complexity.

### Clone tree analysis of intraductal papillary mucinous neoplasms

Fujikura *et al*. performed multi-region whole exome sequencing on 17 patients with intraductal papillary mucinous neoplasms (IPMNs, Fujikura et al. [2020]). Each IPMN contained regions of both low-grade and high-grade dysplasia that were laser capture microdissected, sequenced, and analyzed for evolutionary relationships among samples using the sample tree approach Treeomics [Reiter et al., 2017]. Here we analyzed the clonal evolution of neoplasms from 3 of these patients with PICTograph that illustrate a range of tumor complexity found in real-world applications.

Patient IP29 contained two samples comprised of one low-grade and one high-grade dysplasia with a total of 49 somatic mutations. PICTograph identified 6 mutation clusters and two equally probable clone trees (Figure 5A). Both trees revealed subclone 1 as the truncal clone and branching from the truncal clone into two lineages indicated by subclones 4 and 6. The high-grade dysplasia sample (HG01) is mainly composed of subclones from the lineage beginning with subclone 4, while the low-grade sample (LG02) is composed of the lineage starting with subclone 6 (Figure 5B,C). The difference in the two clone trees lies in the ordering of mutation clusters 2 and 3. In tree 1, cluster 2 is a direct descendant of cluster 3, while in tree 2, cluster 2 and 3 branch from cluster 4 (Figure 5A). Given the available sequencing data, these two possibilities are equally likely.

**Figure 5:**
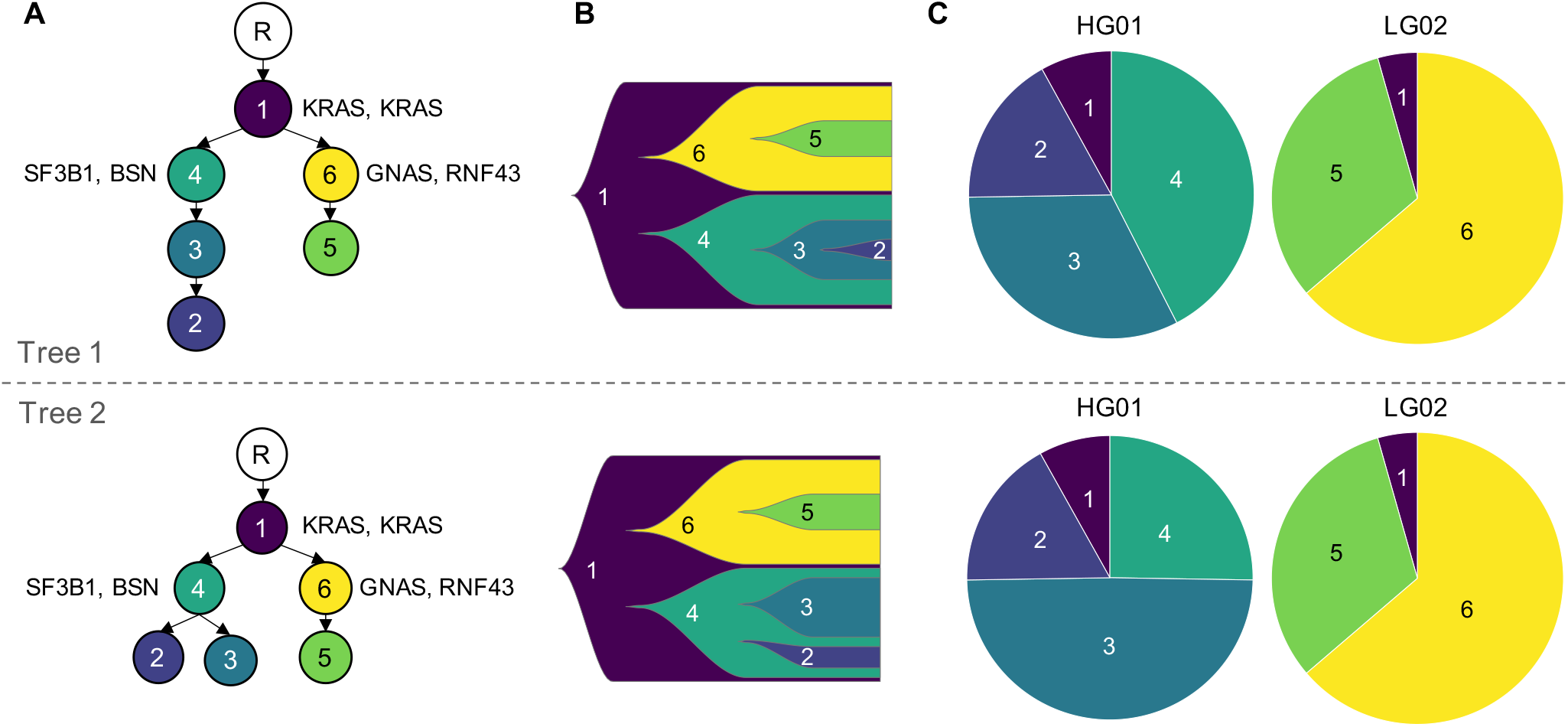
Patient IP29 can be represented by two equally probable clone trees. (A) Clone trees with key driver genes containing mutations labeled next to the clone in which they originated. Clone tree 1 shows a linear evolution of subclones 2 and 3, while clone tree 2 shows branching of subclones 2 and 3. (B) Fish plots created using the average CCF of clones across all samples. (C) Pie charts showing relative proportions of each clone in each sample. The difference in the two clone trees is reflected in the differing proportions of subclones 3 and 4. The linear evolution of subclones 2 and 3 in clone tree 1 means that subclone 2 evolved from a subset of subclone 3, leaving a larger proportion of cells that remained as subclone 4. On the other hand, the branching structure of clone tree 2 means that subclones 2 and 3 evolved from mutually exclusive populations of cells within subclone 4, leaving a smaller proportion of cells that remained as subclone 4.

For Patient IP22, two low-grade and one high-grade dysplasia lesions were sequenced and a total of 83 somatic mutations were detected. PICTograph identified 10 mutation clusters and 6 equally probable clone trees that we summarized by an ensemble tree (Figure 5A). All 6 trees had the same ordering for 8 of the 10 mutation clusters indicated by bold edges in the ensemble (Figure 6A). These analyses revealed that both HG01 and LG05 are comprised of multiple subclones with branches indicating separate lineages starting at subclones 2 and 6, respectively (Figure 6C). Sample LG02 is a mixture of both of these lineages: 75% of one lineage (subclones 2 and 3) and 25% of the other (subclone 6). Distinguishing clonal lineages allows a more granular view of the clonal composition of each sample. In contrast, this subclonal mixing cannot be inferred from the sample tree, which indicates LG02 and HG01 are more closely related to each other than LG05 (Figure 6D). While the clone-tree model places the two *CDKN2A* mutations on distinct branches, the sample tree model indicates a linear evolutionary relationship [Fujikura et al., 2020]. Unlike the sample-tree model, the clone-tree model is consistent with the sample-presence patterns of these two *CDKN2A* mutations that support a branching relationship: one *CDKN2A* mutation is present in LG02 and LG05, while the other is present in LG02 and HG01. Additionally, both *CDKN2A* loci show loss of heterozygosity, suggesting that these two mutations reside in mutually exclusive cell populations. PICTograph identifies mixing of subclones within the sample and identifies a branching structure that is better supported by the data than a linear model.

**Figure 6:**
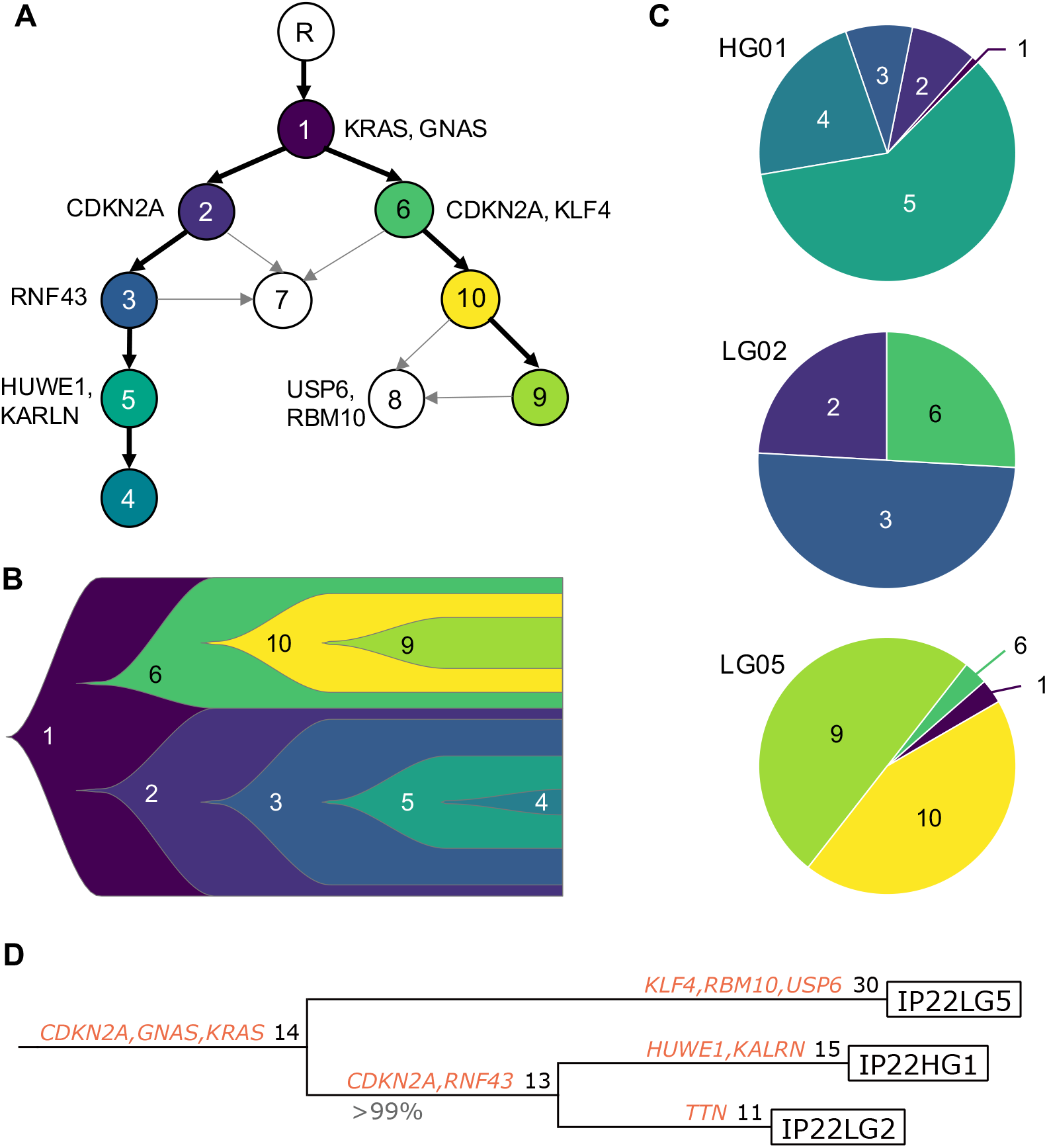
One sample in patient IP22 showed mixing of distinct clonal lineages. (A) Ensemble tree summarizing 6 possible clone trees of equal fitness identified by PICTograph. Key driver genes containing mutations are labeled next to the clone in which they originated. Black edges represent ancestral relationships identified in all 6 trees, while gray edges indicate relationships detected in a subset of the top-scoring trees. (B) Fish plot created using the average CCF of clones across all samples. Only clones with certain immediate ancestors are included and the graphical representation does not convey information about time of origin of subclones as distances between subclones are depicted as equal. (C) Pie charts showing relative proportions of each clone in each sample. The clone tree shows two main lineages that branch from subclone 1. HG01 is composed of subclones in the branch starting with subclone 2, while LG05 contains subclones of the branch starting with subclone 6. LG02 contains a mixture of both of these lineages. (D) The reported sample tree in Fujikura et al. [2020] indicates a single origin for the three regions, with LG2 and HG1 more closely related to each other than LG5.

Two high-grade and two low-grade dysplasia lesions were microdissected from patient IP09 and 155 somatic mutations were identified through whole exome sequencing. PICTograph identified 12 mutation clusters and four equally probable clone trees (Figure 7A). All trees identified subclone 1 containing a driver mutation in *KRAS* as the truncal clone, which then branched into 4 main subclones. Samples HG01, HG03, and LG02 are each comprised of a single lineage, while LG04 is a mix of two lineages (Figure 7C). The diversity of driver mutations in these separate lineages highlight the mutational complexity of pancreatic cancer precursor lesions.

**Figure 7:**
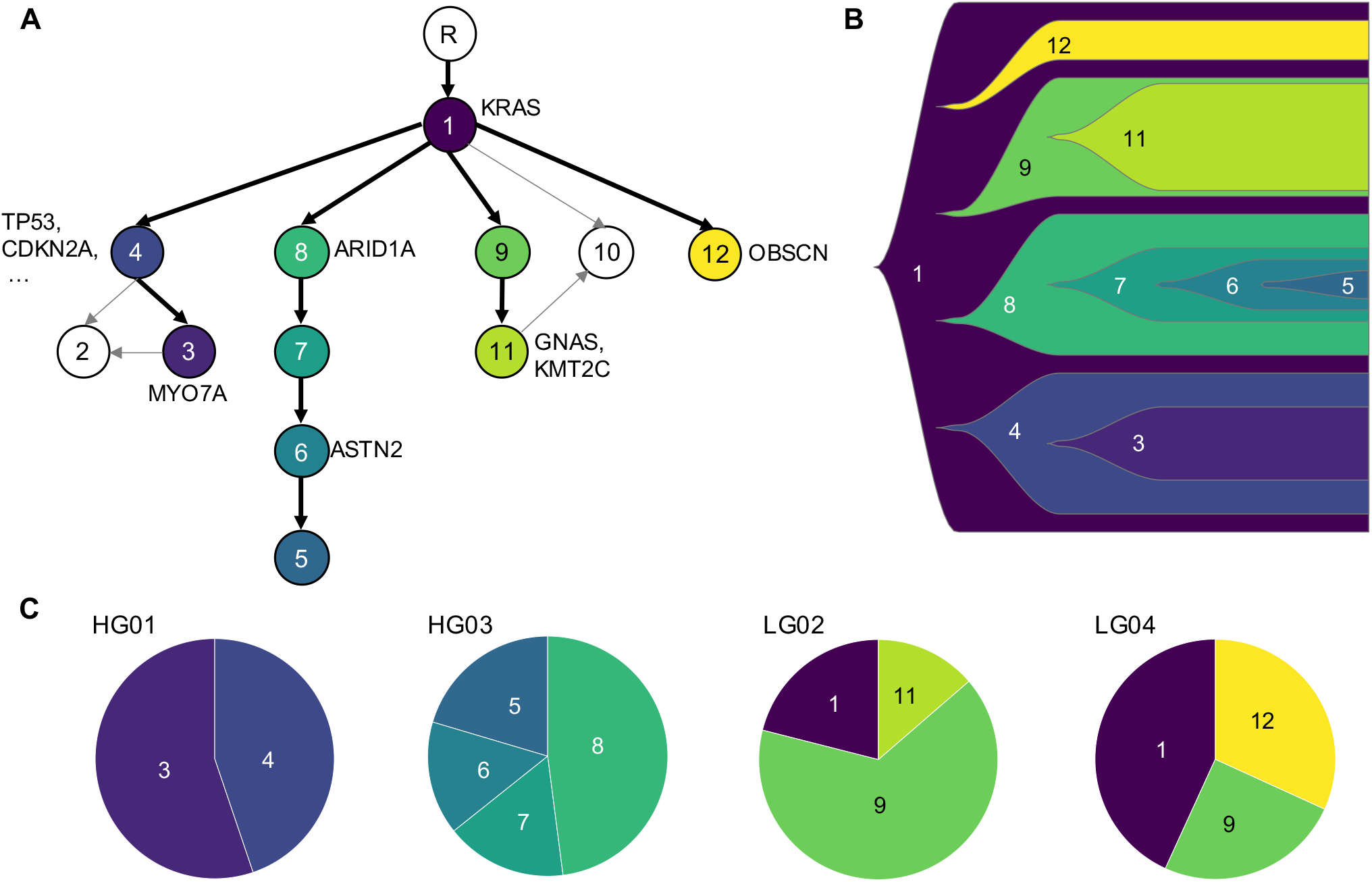
PICTograph identifies four equally probable clone trees for patient IP09. (A) Clone tree with key driver genes containing mutations labeled next to the clone in which they originated. (B) Fish plot created using the average CCF of clones across all samples. (C) Pie charts showing relative proportions of each clone in each sample. The clone tree shows 4 main lineages that branch from subclone 1. Out of the 4 samples, only one samples, LG04, shows mixing of two lineages (subclones 9 and 12). The other three samples contain subclones of a single lineage.

Overall, our analyses captured known early driver mutations in IPMN tumorigenesis and placed known oncogenic driver *KRAS* as the earliest truncal mutation, followed by *GNAS* and tumor suppressor gene alterations in *CDKN2A, RNF43, TP53*, and *ARID1A*. Our analyses also showed mixing of different clonal lineages in 2 of the 3 analyzed cases, highlighting the utility of a clone-tree based approach. Our ensemble-based visualizations of the top-scoring trees revealed consistent evolutionary relationships and different possible evolutionary paths to subclones that are equally likely given the observed data.

## Discussion

We have developed a new algorithm PICTograph that estimates the cancer cell fraction (CCF) of each somatic mutation and infers the most likely evolutionary tree topology from multi-region bulk sequencing data. PICTograph leverages the sample-presence patterns of mutations to inform mutation clustering and to constrain the space of possible trees. By modelling the joint distribution of allelic frequencies across samples, PICTograph resolves cluster memberships and reveals subclonal populations that would otherwise be indistinguishable with single sample analyses. PICTograph performs well over a wide range of simulated multi-sample tumor complexity encountered in experimental applications, and is therefore likely to be of broad utility for a number of cancer types and stages of cancer progression.

While our approach outperforms existing state-of-the-art methods for inferring correct ancestral relationships in a comprehensive series of simulations, PICTograph has several limitations. First, the clustering algorithm assumes that the purity, copy number, and multiplicity of a mutation have already been correctly estimated and, implicitly, that these characteristics are measured without error. Noise of these estimates are not currently reflected in the posterior for the CCFs or in the posterior probability of mutation membership to clusters. Secondly, PICTograph identifies the most probable trees as those that Sum Condition, a criterion based on the infinite sites assumption. Mutations that are lost during evolution would violate this assumption. The critical role of the infinite sites assumption in determining the likely evolutionary relationships is shared by many of the existing methods, including SCHISM, PhyloWGS, PyClone, Canopy, LICHeE. Finally, ancestral relationships for the latent tumor subclones are based on maximum a posteriori estimates of the CCFs and uncertainty of these estimates, while available from the posterior, is not reflected in the set of possible evolutionary trees.

Our application of PICTograph to pancreatic cancer precursor lesions is motivated by previous studies indicating that pancreatic cancers are heterogeneous at earlier stages [Reiter et al., 2017, Fischer et al., 2019, Felsenstein et al., 2018, Wu et al., 2011]. Consistent with these studies, our analyses highlight two patients where different clonal lineages were identified by PICTograph from multi-region sequencing. The absence of mixing of subclonal lineages for the sequenced samples in the third patient may reflect the careful microdissection of the samples sequenced. We anticipate that the application of clone-tree methods such as PICTograph for evolutionary inference will be especially critical for studies employing bulk tissue sequencing where clonal diversity of the bulk sample is likely to be substantial.

## Methods

### PICTograph approach

#### Mutation clustering and cancer cell fraction estimation

As an initial step for CCF estimation, we separated mutations into sets of samples for which the mutation was detected (sample presence) and partitioned the sets into tiers according to the number of samples in the set. For example, a patient with three sequenced samples could have as many as three tiers of mutation sets. Independently for each tier, we evaluated a generative model for the observed number of reads with a somatic mutation *y* for allele *i* in sample *s* that permits inference for the latent CCF:

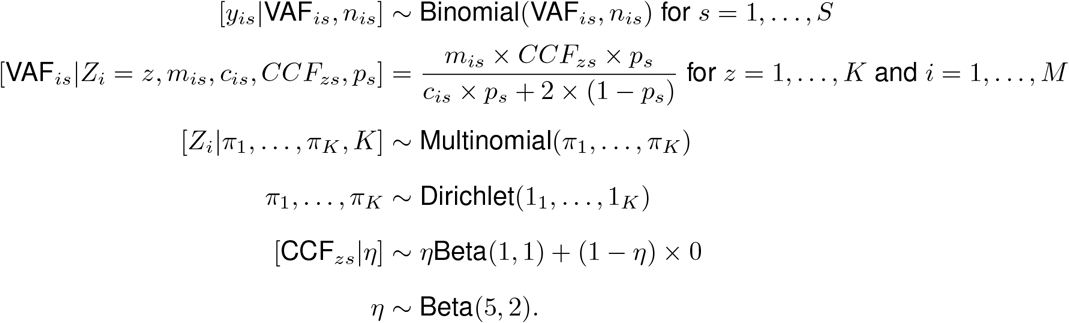

The unobserved parameters *Z*_*i*_ and *CCF*_*zs*_ indicate the cluster membership for *i*th mutation and the cancer cell fraction for cluster *z* in sample *s*, respectively. The joint posterior distribution of {*Z, V AF*, ***π***, *η*} was approximated by Markov chain Monte Carlo (MCMC) implemented using JAGS (version 4.3.0). For each tier, we evaluated a range of possible values for *K* (1 - 15) and selected the *K* that minimized the Bayesian Information Criterion (BIC). With 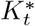 mutation clusters identified in sample presence tier *t*, the total number of mutation clusters for a patient was obtained by 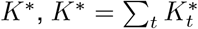.

#### Clone tree inference

PICTograph constructs a rooted mutation tree to represent the tumor clonal evolution, where the root node corresponds to normal cells without somatic mutations and each tree node corresponds to a cluster of somatic mutations with the same cancer cell fraction. A case with *K*^*^ mutation clusters will have trees with *K*^*^ + 1 nodes to include the root node. We define a tumor clone to be the set of mutations along a path from the root to a cluster node. While our Bayesian model provides a joint posterior distribution for {*Z, CCF*}, tree inference is computationally intensive and we limited our analysis to the maximum a posterior estimates of these parameters. Separation of CCF estimation from tree inference has the additional advantage of allowing plug-in estimates for these parameters from other methods.

PICTograph’s tree inference algorithm determines the possible edges between mutation clusters, assembles these directed edges into acyclic graphs (evolutionary trees) consistent with the Sum Condition, and scores trees according to fidelity to lineage precedence and divergence rules. To identify candidate evolutionary relationships between mutation clusters, we identified all directed edges between mutation clusters that were consistent with lineage precedence (3) and sample-presence. A mutation cluster is considered to be present in a sample if its CCF is at least 0.01. Edges consistent with lineage precedence and sample presence have the properties that (i) the descendant cluster must be present in the same or a subset of samples in which the ancestral cluster is present and (ii) the CCF of the descendant cluster are at most 0.1 greater than that of the ancestral cluster in each sample. Conditional on the set of possible directed edges, we applied the Gabow-Myers algorithm [Gabow and Myers, 1978] to enumerate all spanning trees, which are graphs where all the nodes are connected with minimum possible number of edges. Trees identified from this algorithm where the sum of the CCFs of the descendants of a parent node (Sum Condition) exceeded the parent node’s CCF by more than 0.2 were excluded. The 0.1 and 0.2 cutoffs on the lineage precedence and Sum Condition rules, respectively, was motivated by a desire to avoid eliminating trees near the decision boundaries.

All trees in the filtered set were scored by SCHISM [Niknafs et al., 2015] using a fitness function that summarizes violations of lineage precedence by a topology cost and violations of lineage divergence by a mass cost. With *I* → *J* denoting cluster *I* as an ancestor of cluster *J*, the topology cost for the edge connecting two mutation clusters *tc*(*I, J*) is calculated from a binary *Precedence Order Violation* (*POV* matrix), where non-zero entries mark mutation pairs for which the null hypothesis *I* → *J* was rejected. The *Cluster Precedence Order Violation* (*CPOV* matrix) is a straightforward extension of the *POV* matrix in which the hypothesis test is applied to pairs of clusters rather than to pairs of mutations. 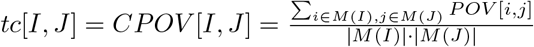 where *M* (*X*) is the set of mutations in cluster *X* and *M* (*X*) denotes the number of mutations in cluster *X*. The topology cost of the tree *TC*(*T*) is then obtained by summing over all the connected clusters in the tree. Lineage divergence assumes that the sum of CCFs of nodes that are descendents, *D*(*n*), of a parent node *p*(*n*) can not exceed the CCF of the parent. While parents with descendant relationships that satisfy lineage divergence have no mass cost, the cost of a violation is given by 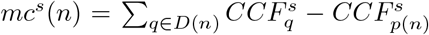. The total mass cost of a tree is obtained as the sum over all nodes in a a tree *N*(*T*), ∑_*n*∈*N*(*T*)_ *mc*(*n*), where *mc*(*n*) is the Euclidean norm of the mass cost across samples, 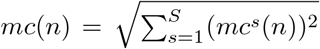. Finally, the fitness of a tree, *F* (*T*), is given by exp(−*f*_*x*_ × [*TC*(*T*) + *MC*(*T*)]). The negative exponent ensures that the highest scoring tree are those with the lowest combined mass and topology costs. In the event of ties for the highest score, PICTograph constructs an ensemble tree with edges weighted by their concordance among constituent trees in the ensemble. Visually, concordance was plotted on a gray scale ranging from black (present in all trees) to light gray (present in a subset of trees).

#### Simulation of CCFs and clone trees

We simulated multi-region (*L* = 3) tumor sequencing data for 100 patients with *K* mutation clusters, setting *K* to 5, 10, or 15 (300 patients total). For each simulated patient, we grew a tree from the root node by sampling the ancestor of each new node from a discrete uniform distribution. For a tree with *k* nodes (including the root), the parent of node *k* + 1 was sampled from the existing *k* nodes each with probability 1*/k*. This process was repeated until the tree had *K* nodes in addition to the root. Next, we simulated *CCF* values for each of the *K* non-root nodes. From the known tree and purity values, we started with the root node with *CCF* = 1 and generated *CCF* values for its direct descendants from a uniform Dirichlet distribution. Moving further along the tree from the root, we continued to simulate *CCF* s for the children of each parent node in a similar manner such that the *CCF* s of a parent node was equal to the sum of the *CCF* s of its children. Next, for each of the *K* nodes, we randomly sampled the number of member mutations ranging from 1 to 10, resulting in a total of *I* mutations where each has a cluster assignment *z*_*i*_, *z*_*i*_ ∈ {1, …, *K*} for *i* ∈ {1, …, *I*}. For each mutation, the multiplicity *m*_*i*_ was randomly assigned to be either 1 or 2 with total copy number *c*_*i*_ equal 2. We calculated the true variant allele frequency as *V AF*_*is*_ = *m*_*is*_*CCF*_*zs*_*/*2. The sequencing depth of each mutation in each sample, *n*_*is*_ was sampled from a Poisson distribution with mean 100 to reflect 100x whole exome sequencing. Finally, the number of alternate reads, *y*_*is*_ was obtained by max(*y*^*^, 1) where *y*^*^ was sampled from a Binomial(*V AF*_*is*_, *n*_*is*_).

#### Implementation of alternative approaches

SCHISM (v1.1.3) was run using the KMeans algorithm for clustering with 10 random initializations each having a minimum and maximum cluster count of 1 and 20, respectively. We used the default genetic algorithm parameters supplied in the SCHISM’s usage example. Canopy was run under settings that do not incorporate copy number alterations as our simulations were limited to single nucleotide variants. For each simulated patient, we ran Canopy with 10 Markov chains using random starts with possible *K*^*^ ranging from 2 to 13, 17, or 20, for true *K* of 5, 10, and 15, respectively. PhyloWGS (v1.0-rc2) was run with recommended default settings and 4 MCMC chains, sciClone (v1.1) was run with minimum depth of 30 and maximum clusters of 20, and PyClone (v0.13.1) was run with default settings. Because LICHeE (v1.0) does not incorporate copy number into its model, we supplied it with cluster CCFs from PyClone and proceeded with default parameters for tree inference. For LICHeE, we limited our analysis to 250 of the 300 simulations as LICHE failed to find valid trees without removing mutation clusters for 50 of the simulations.

#### Assessment of CCF accuracy

Approaches for estimating cancer cell fractions differ according to the mutations assigned to a mutation cluster (membership), the number of mutation clusters identified, and the average cancer cell fraction estimated for each cluster. For a simulated patient with *L* sequenced tumors and *K* clusters, we obtain for each method a *L* by *K*^*^ matrix of estimated cancer cell fractions, ***CCF*** ^*^. To compare the CCF estimates to the true CCFs, we used previously described measures of divergence [Satas and Raphael, 2017]. As these measures are not symmetric (divergence(*x, y*) /= divergence(*y, x*)), we computed divergence in both directions for each method *g* as

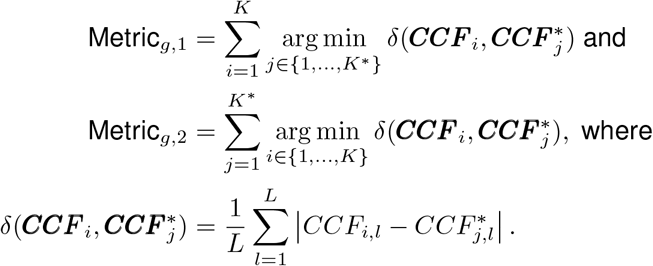

Metric_*g*,1_ provides a distance between estimated and true cancer cell fractions that penalizes underfitting (*K*^*^ too small) but not overfitting (*K*^*^ too large), whereas the opposite is true of Metric_*g*,2_. Therefore, our assessment of accuracy evaluates the joint distribution of these measures of divergence. For each simulation, we can compare two methods *g* and *h* by Δ_*g,h*,1_ = Metric_*g*,1_ − Metric_*h*,1_ and Δ_*g,h*,2_ = Metric_*g*,2_ − Metric_*h*,2_. Omitting the subscripts *g* and *h*, we concisely denote the proportion of simulations for which Δ_1_ and Δ_2_ are positive by *p*_1_ and *p*_2_.

In addition to these divergence measures of CCF accuracy, we compared each method’s ability to recover the true tree by measuring the proportion of mutations that were correctly placed in the best scoring tree generated by each method [Satas and Raphael, 2017]. A mutation pair *m*_1_, *m*_2_ may either have *m*_1_ and *m*_2_ in the same node, or *m*_1_ may be ancestral to *m*_2_, *m*_2_ ancestral to *m*_1_, or *m*_1_ and *m*_2_ may be on distinct branches of the tree. For all pairs of distinct mutations in each sample, we measure whether the reported relationship matched the relationship in the true tree.

#### Analysis of pancreatic cancer precursor lesions

Fujikura et al. [2020] performed whole exome sequencing data analysis for several patients with IPMNs. From each IPMN, 2 to 6 regions were laser capture microdissected. Whole exome sequencing, alignment, and identification of SNVs were obtained as previously described [Fujikura et al., 2020]. Mutations were filtered based on coverage and frequency in the tumor and normal samples, and all non-coding and synonymous variants were removed as well as common germline variants found in databases including dbSNP, the 1000 Genomes Project, Exome Sequencing Project (6500), and Exome Aggregation Consortium (ExAC). Mutations were also validated by visual inspection in IGV. Somatic copy number variants (CNVs) were identified with CNVkit, version 0.9.6 [Talevich et al., 2016] using the matched normal samples obtained from the patients as a reference set. To segment the copy ratio profiles, CNVkit’s default segmentation method and thresholds were used, and samples with >1000 segments were re-segmented using a decreased threshold to reduce the risk of false positive CNV calls. Tumor purity was estimated using somatic mutations in likely copy neutral regions under the assumption that all samples were of at least 40% purity and had multiple clonal somatic mutations. These purity estimates were then used to calculate the integer tumor copy numbers for each segment. Multiplicity, *m*, was estimated by applying constraints such that *m* ≤ *c*_*T*_, with *c*_*T*_ representing tumor copy number. *CCF*s were estimated by PICTograph as previously described. For IP22, we adjusted the estimate of allele-specific copy number for the GNAS mutation from 1 to 2 (out of a total copy number of 3) as a value of 1 for the allele-specific copy number resulted in an extra mutation cluster with CCF values that were incompatible with valid trees.

## Acknowledgments

This work was funded by NIH/NCI CA006973, CA062824, CA12113 (RS); NIH/NCI P50 CA62924, Allegheny Health Network-Johns Hopkins’ Cancer Research Fund, Goldman Pancreatic Cancer Research Center, NIH/NIDDK K08 DK107781 (LDW), and a Johns Hopkins’ Discovery Award (RK and LDW).

